# Novel ACE2 nanoparticles universally block SARS-CoV-2 variants in the human respiratory tract

**DOI:** 10.1101/2022.05.05.490805

**Authors:** Cécile Sauvanet, Moara Lemos, Armel Bezault, Borja Rodríguez de Francisco, Michael CW Chan, Kenrie PY Hui, Ka-chun Ng, John M Nicholls, Niels Volkmann, Dorit Hanein

## Abstract

The continual evolution of SARS-CoV-2 has challenged the efficacy of many COVID19 vaccines and treatment options. One strategy that evades viral escape is using the entry receptor, human Angiotensin-Converting Enzyme 2 (hACE2). Soluble hACE2 receptor domains show potential as decoys but genetic modifications are necessary to provide sufficient efficacy. However, these engineered constructs are potentially susceptible to viral escape. We combined native hACE2 with viral vectors to form nanoparticles presenting hACE2 analogous to human cells. Cell-based viral infection assays and cryogenic *in-situ* tomography show that hACE2 nanoparticles sequester viruses through aggregation, efficiently blocking entry of SARS-CoV-2 and its variants in model cell systems and human respiratory tract explants using native hACE2. Thus, we show that hACE2 nanoparticles have high potential as pan-variant COVID19 therapeutics.

---

The COVID-19 pandemic is wreaking havoc on health and economy on a global scale. The disease is caused by infections with Severe Acute Respiratory Syndrome Coronavirus 2 (SARS-CoV-2) (*1*). The virus enters human cells via interactions between its spike protein and the human angiotensin-converting enzyme 2 (hACE2) (*2*). The virus can enter through membrane fusion, which also requires the presence of Transmembrane Serine Protease 2 (TMPRSS2) (*3*), or through the clathrin-mediated endocytic pathway (*4*). In either case, the presence of hACE2 on the cell surface is obligatory for viral entry.

Several vaccines and treatment approaches have been developed against SARS-CoV-2. Most of these options rely on the actions of antibodies that bind to exposed viral proteins and prevent cell entry. A major concern with this strategy has been the appearance of different naturally evolving variants of SARS-CoV-2 that cause not only enhanced transmissibility but also tend to decrease susceptibility to antibody neutralization or therapeutics. The delta and omicron variants are particularly problematic in this regard. The delta variant has caused deadly second waves of disease around the world (*5, 6*) and is known to have a high level of transmissibility and virulence when compared to the wild-type virus. The omicron variant carries 15 mutations in the ACE2 receptor binding domain alone (*7*), which significantly weakens the efficiency of vaccines (*8*) and antibody treatments (*9*) with the BA.2 sublineage escaping nearly all monoclonal antibodies currently authorized for therapeutic treatment (*10*). The omicron variant also exhibits increased escape from antibodies generated by previous infections or vaccinations, causing numerous reinfections and breakthrough infections. In addition, treatment with monoclonal antibodies has been linked to rapid viral evolution and the appearance of escape mutations (*11, 12*), suggesting that antibody-based treatments may promote the appearance of new variants of concern.

Treatment options that do not depend on the action of neutralizing antibodies include those that target the viral replication process where the drug interferes by binding to the viral protease or to the viral polymerase (*13*). Because these inhibition mechanisms involve binding to viral proteins, which can be subject to mutations, this strategy is also prone to be affected by viral escape. In addition, drugs that are targeting the replication process are potentially more likely to cause side effects because they act inside the cell rather than preventing viral entry altogether. For example, the protease inhibitor Molnupiravir has raised concerns about potential long-term side effects including cancer and birth defects (*14*). COVID19 is likely to remain a leading cause of death in the future and, based on current experience, new variants will continue to emerge. Thus, there is a continuing need for effective therapeutics, especially as new variants are expected to escape natural and vaccine-induced antibodies at least to some extent.

One viable strategy to combat viral escape is to make use of the obligatory entry receptor, in this case hACE2, to prevent viral entry. Even heavily mutated viruses must preserve their ability to bind to this receptor in order to enter their target cells. Presenting soluble hACE2 constructs (sACE2) outside the cell engage the wild-type virus as decoys, thereby diminishing its cell entry capacity (*15*). In particular, sACE2 constructs engineered for exceptionally high affinity, which rivals the affinity of monoclonal antibodies, were shown to bind efficiently to several SARS-CoV-2 variants (*16*). One of these high-affinity decoys was shown to be effective against SARS-CoV-2 variants when administered intravenously to humanized mice (*17*) and inhalation of the same construct was found to increase survival in humanized mice inoculated with a lethal dose of virus (*18*). However, the mutations in the spike/hACE2 interface of these engineered sACE2 constructs, necessary to boost the efficacy of the decoy activity, make the construct potentially prone to viral immune escape.

Here, we targeted SARS-CoV-2 cell entry by constructing an efficient decoy that involves a carrier system based on murine leukemia virus (MLV), which has been extensively used as a viral vector in gene therapy studies (*19*), studded with full-length, native hACE2 on its surface (hACE2 nanoparticles). To monitor the effect of these hACE2 nanoparticles on SARS-CoV-2 entry using luciferase activity assays (*20*), we also generated MLVs carrying luciferase mRNA (control MLVs) and pseudoviruses (*21*) composed of control MLVs pseudotyped with wild-type SARS-CoV-2 spike, delta-variant spike, or omicron-variant spike.

Following pseudovirus infection, we used cellular cryogenic electron tomography (cryo-ET) to verify the morphology of pseudoviruses and to independently confirm infection of Vero-E6 African green monkey kidney cells. Vero-E6 cells are known to support SARS-CoV-2 replication and were shown to report accurately on SARS-CoV-2 pseudovirus infection rates (*22*). Vero-E6 cells endogenously express African green monkey ACE2, which shares 100% sequence identity with hACE2. The presence of SARS-CoV-2 spikes in both the pre- and post-fusion configurations protruding from the pseudovirus surface are clearly visible in the cryo-ET data and the structural signature of the pseudovirus capsid is highly distinct, facilitating unanimous identification of the pseudovirus not only outside but also within cells (Fig. S1). Our analysis of pseudoviruses inside Vero-E6 cells revealed that pseudovirus cell entry in these cells is accomplished via the clathrin-mediated endocytic pathway (Fig. S1). This mechanism is consistent with entry of SARS-CoV-2 in the absence of TMPRSS2 (*3*) and is the preferred pathway of entry for the omicron variant (*8*).

We verified that hACE2 is required for entry of pseudoviruses using luciferase activity assays using the Baby Hamster Kidney fibroblasts 21 (BHK-21) cell system (*22*). We used parental BHK-21 cells, which lack hACE2, and BHK-21 cells transiently expressing hACE2 to verify that pseudoviruses only enter cells that carry hACE2 (Fig. 1A). To quantify the effect of hACE2 nanoparticles on pseudovirus cell entry we conducted luciferase activity assays with the Vero-E6 system. We found that the cell entry signal from SARS-CoV-2 pseudoviruses in the presence of hACE2 nanoparticles is statistically indistinguishable from the signal from the pseudovirus-free Control for wild-type, delta-variant, and omicron-variant pseudoviruses alike (Fig. 1B and C), This finding indicates that hACE2 nanopraticles are completely abolishing SARS-CoV-2 cell entry. Quantification of the hACE2 concentration on the nanoparticles (Fig. 1D and E) showed that, in the concentration used for our experiments, it corresponds to about 14 nM of hACE2 receptor binding domain. To achieve a similar effect on entry suppression using sACE2 alone, the sACE2 concentration needs to be increased to at least 500 nM.

**Fig. 1.**
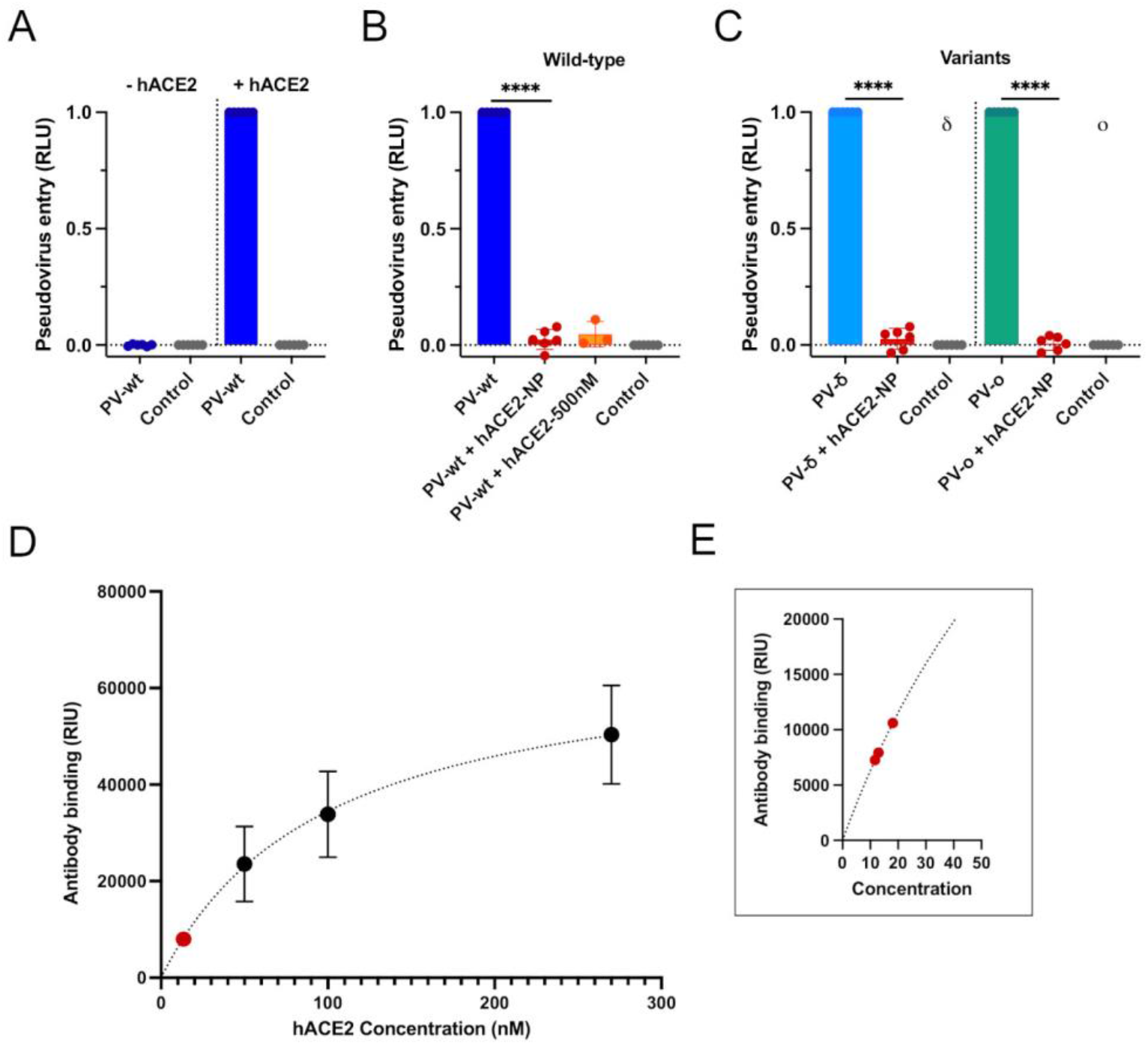
hACE2 nanoparticles block SARS-CoV-2 pseudovirus entry. **(A)** Luciferase activity assays were performed after overnight incubation of parental BHK-21 cells (-hACE2) or hACE2-expressing BHK-21 cells (+hACE2) with wild-type pseudoviruses (PV-wt). SARS-CoV-2 pseudoviruses enter cells and generate a luciferase signal only for cells expressing hACE2. All luciferase data were normalized to correspond to zero for the signal from control MLVs only carrying luciferase mRNA (Control) and set to one for the signal from pseudovirus only. The bars represent the mean ± standard deviation of experimental replicates. **(B)** Luciferase activity assays of Vero-E6 cells with wild-type pseudovirus (PV-wt) in the absence and presence of hACE2 nanoparticles (hACE2-NP) or 500nM soluble hACE2 (hACE2-500nM). **(C)** Luciferase activity assays of Vero-E6 cells with delta-variant (PV-δ) or omicron-variant (PV-o) pseudovirus in the absence and presence of hACE2 nanoparticles (hACE2-NP). Entry of pseudoviruses into Vero-E6 cells is blocked by hACE2 nanoparticles for all variants including wild type. **(D)** Quantification of hACE2 level in hACE2 nanoparticles by western blot (Fig. S2). The curve plots the intensity of the signal from hACE2 antibodies versus the corresponding concentration of soluble hACE2. The red dot marks the average location of the hACE2 nanoparticle intensities. **(E)** Intensities from three independent samples of hACE2 nanoparticles overlaid on the calibration curve were used to extrapolate hACE2 concentration.

To investigate if our results translate to human SARS-CoV-2 infection, we chose tissue explants of the human respiratory tract as a model system. Because mice have mismatched ACE2 receptors, even genetically engineered, humanized mice are not a perfect substitute for human infection (*23*). In contrast, while *ex vivo* cultures of human respiratory tract lack *in vivo* effects of the adaptive immune and systemic inflammatory responses, they retain cytoarchitecture and three-dimensional organization of the tissue, including cell-cell and cell-matrix interactions. As such, these *ex vivo* cultures provide one of the most useful models to compare viral replication kinetics, cell tropism, and innate immune responses of SARS-CoV-2 in the human respiratory tract (*24, 25*). We thus tested the effect of hACE2 nanoparticles on the viral replication kinetics of SARS-CoV-2 delta variant in *ex vivo* cultures of human bronchus tissues (*26*). We chose the delta variant for these experiments because it caused the most deadly wave of the disease and is thus the most clinically significant SARS-CoV-2 variant so far (*5, 6*). We found that the presence of hACE2 nanoparticles abolishes viral replication down to the level of detectability (Fig. 2).

**Fig. 2.**
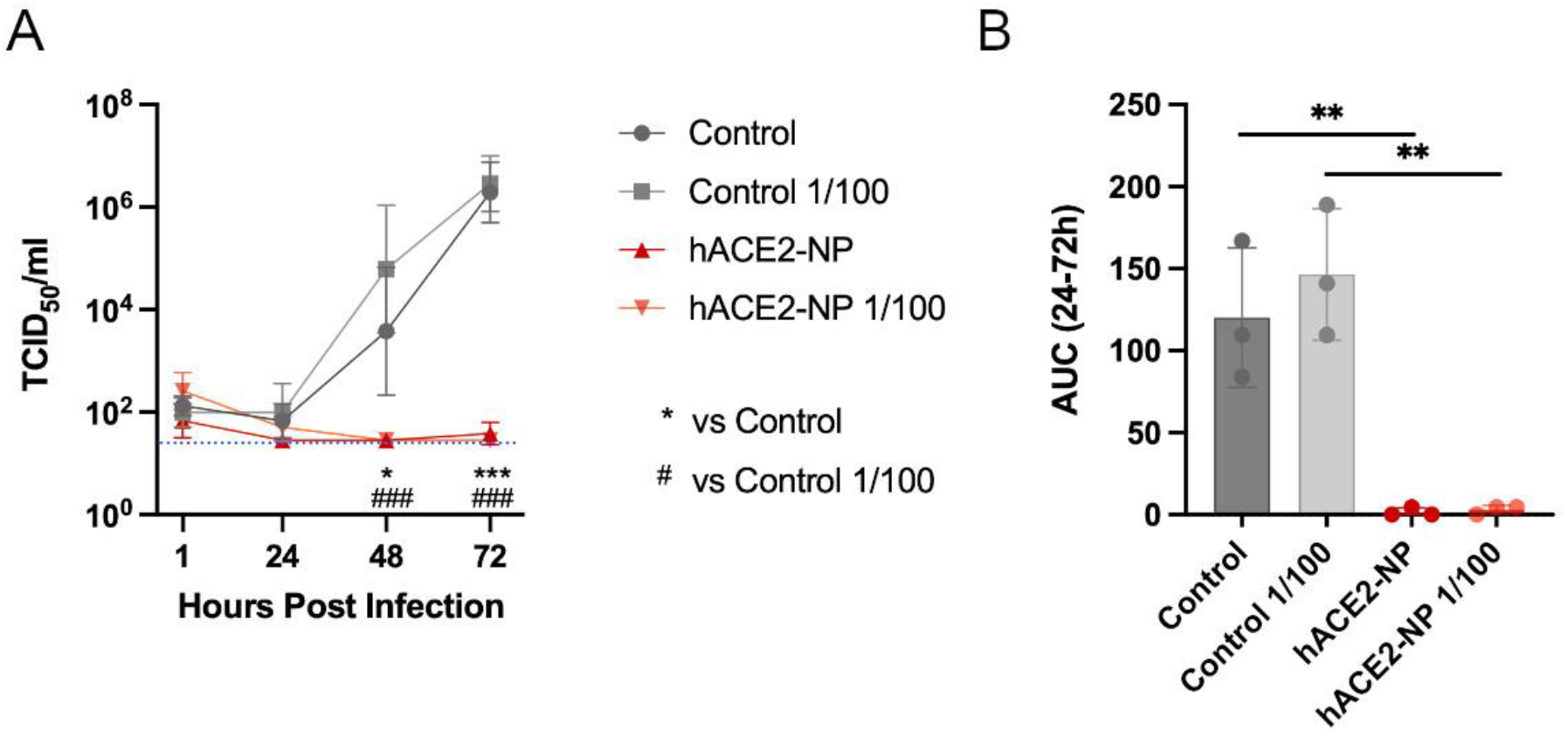
hACE2 nanoparticles block SARS-CoV-2 virus entry in *ex vivo* human bronchus explants. Effect of hACE2 nanoparticles on the viral replication kinetics of SARS-CoV-2 delta variant in *ex-vivo* cultures of human bronchus. Human *ex-vivo* explant cultures of bronchus were pre-treated with control MLVs (Control) or hACE2 nanoparticles (hACE2-NP; neat or 1/100 diluted) and then infected with delta-variant SARS-CoV-2 viruses at 37°C. Virus released in the culture supernatants were measured over time by TCID_50_ assays. **(A)** Viral replication kinetics of Delta in human *ex vivo* cultures of bronchus. The horizontal blue dotted line denotes the limit of detection in the TCID_50_ assay. Data are presented as geometric mean values (n=3) (±SD). Statistics were performed using Two-way ANOVA followed by Tukey’s test. *p<0.05, ***p<0.001 vs control MLVs; ###p<0.001 vs control MLVs 1/100. **(B)** Viral titers are depicted as area under the curve (AUC). Bar-charts show the mean (n=3) (±SD). Statistics were performed using One-way ANOVA followed by a Tukey’s multiple-comparison test. **p<0.01.

To elucidate the mechanism of hACE2 nanoparticles entry inhibition, we employed cryo-ET to characterize the hACE2 nanoparticles and their interactions with SARS-CoV-2 pseudoviruses. Structural analysis of the hACE2 nanoparticles revealed a high density of particles protruding from the MLV membrane that are fully consistent with the structure of membrane-embedded ACE2 dimers (*27*) (Fig. 3A and B). hACE2 nanoparticles engage with multiple SARS-CoV-2 pseudoviruses leading to the formation of large aggregates, efficiently sequestering pseudoviruses before cell entry (Fig. 3C and D).

**Fig. 3.**
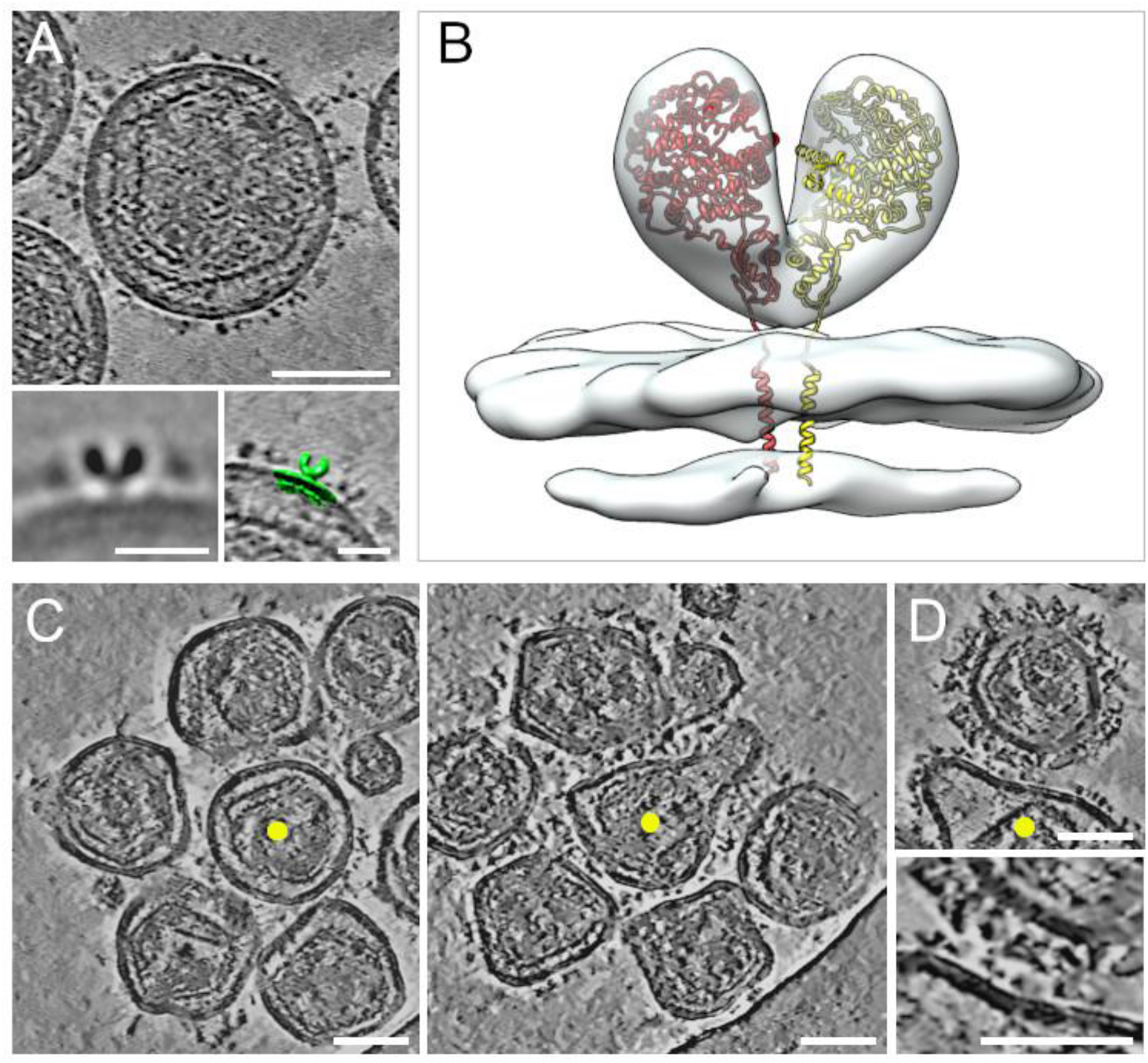
Cryogenic electron tomography of hACE2 nanoparticles. All virtual slices in this figure are 12-nm thick. Bars in main panels = 50 nm; bars in insets = 20 nm. **(A)** Slice through a tomogram of an hACE2 nanoparticle showing dense decoration with hACE2 dimers. Insets show two-dimensional average of the protein density protruding from the nanoparticles (left) and overlay with surface representation (green) of three-dimensional (3D) reconstruction of membrane-bound particles (right). **(B)** Surface representation of 3D reconstruction obtained by sub-tomogram averaging of membrane-bound particles (Fig. S3). The atomic structure of the ACE2 dimer (monomers in yellow and red), extracted from the open conformation of the hACE2-B0AT1 complex (PDB: 6MD1), was fitted into the density showing a good match. **(C)** Virtual slices through tomograms of hACE2 nanoparticles (marked by yellow dots) interacting with several SARS-CoV-2 pseudoviruses at the same time. **(D)** hACE2 nanoparticle (yellow dot) interacting with multiple spikes on a SARS-CoV-2 pseudovirus. The lower panel shows an enlarged view of the interaction region.

We here provide evidence that hACE2 nanoparticles are highly efficient in preventing infection of Vero-E6 cells by SARS-CoV-2 pseudoviruses as well as infection by live SARS-CoV-2 delta variant viruses of *ex vivo* human bronchus tissue. Our results indicate that the blockage of virus entry occurs not only via competition of membrane-embedded hACE2 receptors in the nanoparticles with hACE2 cell receptors (decoy effect) but also via aggregation through the engagement of hACE2 receptors on the same nanoparticle with spike proteins of multiple SARS-CoV-2 viruses. Our studies suggest that hACE2 nanoparticles provide a potent and effective viral entry neutralization effect and thus present a promising opportunity to develop new COVID19 treatment options that are less prone to viral escape than current strategies based on antibodies or engineered sACE2 decoys. The *ex-vivo* results show that the inhibition by hACE2 nanoparticles remains highly efficient in human respiratory tracts even if the concentration is reduced by two orders of magnitude. Soluble hACE2 constructs have been shown to be well tolerated in clinical trials (*28*) and MLV-based particles have a long history of clinical applications in the context of viral gene therapy (*29*). It was shown that engineered sACE2 decoys can be administered efficiently using aerosols (*18*). With hACE2 nanoparticles matching the size of the virus, it is expected that administration with aerosols will efficiently localize the nanoparticles in the same locations within the respiratory tract where the virus ends up. Our studies, combined with these practical facts can accelerate the development of preventative as well as early stage COVID19 therapeutics based on hACE2 nanoparticles.

## Materials and Methods

### Cell lines

Human primary embryonic kidney 293T/17 (HEK293T/17) cells, monkey kidney Vero E6 cells (Vero-E6), and Baby hamster kidney fibroblast 21 (BHK-21) cells were obtained from the American Type Culture Collection (ATCC® CRL-11268, ATCC® CRL-1586™, ATCC® CCL-10™). HEK293T/17 cells were maintained in a humidified 5% CO_2_ incubator in complete Dulbecco’s Modified Eagle Medium (DMEM) supplemented with 100 U/ml penicillin, 100 μg/ml streptomycin, 10 mM 4-(2-hydroxyethyl)-1-piperazineethanesulfonic acid (HEPES), and 10% fetal bovine serum (FBS) (Sigma-Aldrich). HEK293T/17 cells were used to produce pseudoviruses with spike protein from wild-type SARS-CoV-2, delta variant, and omicron variant. HEK293T/17 cells were also used to produce MLV control and hACE2 nanoparticles. Vero E6 and BHK-21 were maintained at 37°C with 5% CO_2_ incubator in Modified Eagle Medium (MEM) supplemented with 1X non-essential amino acid, 2mM L-glutamine, 100 U/ml penicillin, 100 μg/ml streptomycin, and 10% FBS (all from Sigma-Aldrich). All cell lines were tested for mycoplasma every three months using the Lookout Mycoplasma PCR detection kit (Sigma-Aldrich) and were authenticated by ATCC®.

### Plasmids

The plasmids for MLV production, pCMV-MLV-gag-pol and pTG-Luc, were kindly provided by Jean Dubuisson (Institut Pasteur de Lille). pcDNA3.1-SARS2-Spike (*2*) and pcDNA3.1-hACE2 (*2*) were obtained from Addgene (Addgene plasmids #145032; #145033); pCI-SARS2-Spike expressing a codon optimized version of the gene coding for S protein of SARS-CoV-2 was provided by Nicolas Escriou (Institut Pasteur). pUNO1-SpikeV8 carrying delta variant spike and pUNO1-SpikeV11 carrying omicron (B.1.1.529/BA.1 lineage) spike were purchased from InvivoGen (catalogue numbers p1-spike-v8 and p1-spike-v11 respectively).

### Pseudovirus and hACE2 nanoparticles production

Control MLVs, MLV-based pseudoviruses carrying SARS-CoV-2 wild-type, delta, or omicron spike protein, and MLV-based hACE2 nanoparticles were prepared as previously described (*21*). In brief, HEK-293T/17 cells were co-transfected using Lipofectamine 2000 (Life Technologies) with MLV Gag-Pol packaging construct and the MLV transfer vector encoding a luciferase gene reporter either by itself (MLV-control) or with one of the Spike protein or hACE2 protein expression vectors for 48h at 37°C or 72h at 33°C. The resulting particles were released into the supernatant, which was then harvested and filtered through 0.45µm membranes, aliquoted and stored at −80 °C.

### Pseudovirus cell entry assays

Vero-E6 and BHK-21 cells were plated into 24 wells at a dilution of 5 × 10^5^ cells/ml. One day prior, 60 mm dishes of BHK-21 cells were transiently transfected with 2.5 μg human ACE2 or no DNA for control using standard lipofectamine 2000 (Life Technologies) protocols. After 24hrs, 200 µl of pseudovirus were added to the wells, after washing 3X in DPBS, incubated for 1h at 37°C in 5% CO_2_ incubator before adding 300µl of complete MEM over a day (7hrs) or for overnight incubation. Cells were washed in MEM without serum, and 100 µl One-Glo-EX (Promega) was added to the cells in equivalent MEM with no serum volume and incubated in the dark for 5 min. 180µl of the mixture was moved to 96 well plates prior to reading. For the competition assays, experiments were performed in 96 wells plates. Prior to cell infection, 40 μL of hACE2 nanoparticles or the desired concentration of soluble hACE2 (provided by Ignacio Fernández and Félix Rey, Institut Pasteur) were added to the cell culture and incubated for 30 min before addition of pseudoviruses (40 µl). After an additional 30 min of incubation, full media was added to pursue overnight incubation. The luciferase signal was determined in a Varioskan LUX microplate reader (Thermo Fisher Scientific) at 10 readings for each well at intervals of 30 s. Measurements were done in triplicate. A slight increase of luciferase activity was observed after 7hrs of interaction but to observe the potent effect of hACE2 nanoparticles overnight incubation was chosen as reference point. Relative luciferase units (RLU) data were normalized to pseudovirus signal set to one and the signal from controls without pseudovirus set to zero. All normalized data were then statistically analyzed and represented using GraphPad Prism, version 9.3.1 (GraphPad Software).

### Western Blotting

hACE2 nanoparticles and soluble hACE2 were thawed and 4X SDS loading buffer was added prior to boiling for 5 min at 95°C. The same quantity of samples used for the competition assay was run on a 4%–20% gradient Tris-Glycine Gel (BioRad) and transferred to nitrocellulose membrane. A Goat anti-hACE2 polyclonal primary antibody (at 1:100 dilution corresponding to a concentration of 2 µg/ml; Bio-techne, catalogue number #AF933) and an Alexa Fluor 488-conjugated donkey anti-goat secondary antibody (1:10,000 dilution, Jackson Laboratory, Catalog number # 305-547-003) were used for Western blotting. Western blots were imaged using a BioRad ChemiDoc MP Imaging System.

### SARS-CoV-2 delta variant isolation

Delta (B.1.617.2, hCoV-19/Hong Kong/CM21000383_WHP4738/2021) variant was isolated and propagated in Vero-E6-TMPRSS2 overexpressing cells (*30*). The cell line was cultured in DMEM with 10% FBS. The original clinical specimen was collected from SARS-CoV-2 confirmed patients in Hong Kong in 2021 and isolated as previously described (*24*). The virus stock was aliquoted and stored frozen at −80°C. Aliquots were titrated to determine plaque forming units. The experiments were carried out in a Bio-safety level 3 (BSL-3) facility at the School of Public Health, LKS Faculty of Medicine, The University of Hong Kong.

### Ex-vivo cultures and infection of human bronchus tissues

Fresh non-tumor bronchus tissues were obtained from patients undergoing elective surgery in the Department of Surgery at Queen Mary Hospital (Pok Fu Lam, Hong Kong, China) and were removed as part of routine clinical care but surplus for routine diagnostic requirements as detailed previously (*26, 31*). The virus infection procedures were performed as previously described (*26*). Briefly, similar sized pieces of human bronchus tissues were treated with MLV-control or hACE2 nanoparticles for 30 min at 37°C and then infected with SARS-CoV-2 at 4×10^4^ plaque forming units for 1 h at 37°C. Each tissue fragment was washed three times to remove residual virus inoculum and incubated at 37°C with fresh medium. Aliquots of culture medium were removed at times indicated and stored at −80°C until titration. Infectious viral titers in culture supernatants were assayed by TCID_50_ in Vero-E6-TMPRSS2 cells. Informed consent for the use of human tissues was obtained from all subjects and approval was granted by the Institutional Review Board (IRB) of the University of Hong Kong and the Hospital Authority (Hong Kong West) (IRB approval no: UW 20-862).

### Viral titration by TCID_50_ assay

Vero-E6-TMPRSS2 cells were seeded in 96-well tissue culture plates one day before the virus titration (TCID_50_) assay. Cells were washed once with PBS and DMEM (Gibco) with 2% fetal bovine serum (Gibco) supplemented with 100 units/ml penicillin and 100 µg/ml streptomycin (Gibco) was added. Serial dilutions of virus supernatant, from 0.5 log to 7 log, were performed and each virus dilution was added to the plates in quadruplicate. The plates were observed for cytopathic effect daily. The end point of viral dilution leading to CPE in 50% of inoculated wells was estimated using the Kärber method (3*2*). Area under the curve (AUC) was calculated from the viral titers from different time points indicated on the Y-axis.

### Quantification and Statistical Analysis

For western blots and luciferase activity assays, experiments were performed multiple times with at least two independent preparations of each sample. Quantification of the intensity of western blot bands was realized by densitometric analysis using Fiji (*33*), version 2.1.0. Soluble hACE2 samples of different concentrations were used to establish a standard curve, which was used to determine hACE2 level on the nanoparticles. Luciferase data from individual experiments were background corrected by subtracting the signal from control MLVs and then normalized by setting the background corrected signal from unperturbed pseudovirus to one. One-sided, one-tailed t-tests were performed against 1 and 0 for all other normalized, background-corrected conditions; two-tailed t-tests were used to compare the normalized, background corrected signals from pseudovirus in the presence of hACE2 nanoparticles (hACE2-NP) or soluble hACE2.

Significance of differences was determined after Bonferroni correction. The difference between pseudovirus alone and pseudovirus in the presence of hACE2-NP was statistically highly significant (wild-type p-value: 1.73 e^-8^; delta p-value: 2.18 e^-8^; omega p-value: 2.66 e^-9^) and there was no statistically significant difference with Control (wild-type p-value: 0.11; delta p-value: 0.10; omega p-value: 0.30). There was no statistically significant difference between wild-type pseudovirus in the presence of hACE2-NP and 500nM soluble hACE2 (p-value: 0.50). The difference between parental BHK-21 cells in the presence and absence of wild-type pseudovirus was not significant (p-value: 0.37) and the difference between parental and hACE2-expressing BHK-21 cells in the presence of wild-type pseudovirus was highly significant (p-value: 1.57 e^-13^). Statistics for the TCID_50_ assays were performed using two-way ANOVA followed by Tukey’s test. Statistics for the AUC analysis was performed using One-way ANOVA followed by a Tukey’s multiple-comparison test.

### Cryo-ET sample preparation of MLV-based samples

Pseudovirus samples were concentrated 10 to 20 times by ultracentrifugation (Optima MAX-XP Ultracentrifuge; Beckman Coulter) for 30 min, at 4°C in a TLA-55 rotor (Beckman Coulter) at maximum speed (186000 g) then resuspended in similar media overnight at 4°C. Samples were fixed in 4% formaldehyde (Tousimis Research Corporation; catalogue number #1008A) diluted in full media for 30 min at 37°C then stored at 4°C. For vitrification, samples were loaded onto plasma cleaned (Solarus II Plasma Cleaner; Gatan, Inc.) Holey Carbon Film R 1.2/1.3 Copper 400 mesh EM grids (Quantifoil Micro Tools GmbH; catalogue number #N1-C14nCu40-01), incubated for 30 sec prior to plunge freezing in liquid ethane either using a manual plunger or a Vitrobot Mark IV system (Thermo Fisher Scientific). For mixtures between pseudoviruses and hACE2 nanoparticles, the samples were resuspended in similar media for 2-3 hrs. Mixtures of pseudovirus and hACE2 nanoparticles (1:1 volume) were incubated for 30 min at 37°C prior to vitrification. For all samples, ∼1 µl of 5 nm protein A Gold (UMC Utrecht) were added as fiducial markers to assist with tilt series alignment.

### Cryo-ET data collection of MLV-based samples

Data were acquired on a Titan Krios G3i (Thermo Fisher Scientific) operated at 300kV in parallel beam condition, a K3 Summit direct electron detection device and a BioQuantum energy filter (Gatan Inc.) operated in zero-loss mode with a slit width of 20 eV. Automated data collection was carried out using SerialEM (*34*), version 3.8, at a nominal magnification of 42K x with a calibrated pixel size of 0.21 nm. Tilt series were acquired following the dose symmetric scheme (*35*) between +60° and −60° with a step size of 3°. The dose rate was adjusted to 23 counts/pixel/s resulting in a total dose of 100e^-^/Å² per tilt series, and each tilt image was acquired in electron counting mode and fractionated in 13 to 14 frames. The defocus range was −5 to −8 μm.

### Cryo-ET sample preparation of pseudovirus-infected cells

Vero-E6 were seeded on holey carbon film R 5/20 gold 200 mesh EM grids (Quantifoil Micro Tools GmbH; catalogue number # N1-C43cAuE1-01) at a cell density of 5000 cells per grid, incubated in complete media up to 24 hrs at 37°C in 5% CO_2_ incubator. Pseudoviruses were then incubated with the cells from 30 min to overnight. Cells were washed in MEM without serum and then fixed in a structure-preserving buffer (*36*) (0.1 M Pipes, 1 mM EGTA, and 1 mM MgSO_4_) containing formaldehyde 4% for 30 min at 37°C. Plasma membranes of cells were stained for light microscopy using WGA 647 according to supplier protocol (Invitrogen, Thermo Fisher Scientific). Grids were then stored at 4°C in PHEM buffer and 1% formaldehyde. Next, the grids were screened for cell density and integrity on a DMi8 light microscope (Leica Microsystems). Samples that passed the screening were vitrified in liquid-nitrogen cooled liquefied ethane using either a manual plunger, Vitrobot Mark IV system (Thermo Fisher Scientific) or EM GP Model 1 plunge freezer (Leica Microsystems). For all samples, 1µl of 10 nm protein A Gold (UMC Utrecht) was added as fiducial markers to assist with tilt series alignment. The vitrified grids were then mounted into grid support rings (Auto-grid TEM sample holder, Thermo Fisher Scientific), and transferred to a cryogenic light microscopy (cryo-LM) shuttle for screening on a DM6 cryogenic light microscope (Leica Microsystems) to verify the integrity of the samples following vitrification.

### Cryo-FIB Lamella preparation of pseudovirus-infected cells

After cryo-LM screening, autogrids were transferred to an Aquilos-2 cryogenic focused ion beam scanning electron microscope (cryo-FIB-SEM) (Thermo Fisher Scientific). Lamella sites were chosen based on the cell membrane signal obtained from the DM6 cryo-Light microscope (Leica Microsystems) correlated with the SEM images using MAPS 3.14 software, version 3.14.11.45 (Thermo Fisher Scientific). A total of 30 cells incubated with pseudovirus were prepared using either the manual interface (XT User: v20.1.0 - 20.1.0.7012) or semi-automated software (AutoTEM 2.2.0 core 10.0.3) (Thermo Fisher Scientific) in the Aquilos-2 cryo-FIB-SEM. Briefly, cells were coated with an organometallic platinum layer for 12 s and gradually thinned in 4 steps at a stage angle of 15–18° using a Ga^+^ beam to yield lamellae with 200-300 nm thickness after the final milling step. Micro-expansion joints were used to improve lamella stability. Progress was monitored by SEM imaging at 2–10 kV with an Everhart-Thornley Detector (ETD) detector. The final polishing step was performed manually.

### Cryo-ET data collection of pseudovirus-infected cells

For pseudovirus-infected cells and lamellae, cryo-ET data were collected either using SerialEM on a Titan Krios G3i (Thermo Fisher Scientific) operated at 300kV, equipped with a K3 direct electron detector with a BioQuantum energy filter (Gatan Inc.) with energy slit set to 20 eV or a Glacios microscope (Thermo Fisher Scientific) operated at 200kV, equipped with a Falcon 3EC direct electron detector (Thermo Fisher Scientific). The autogrids were rotated 90° between Aquilos-2 and the respective electron microscope to optimize the relationship between the tilt-axis and the lamella orientation. Both instruments were operated in parallel beam condition. For each grid, an overview image (atlas) of the entire grid was acquired to locate lamellae or pre-screened cells for direct imaging. Markings on the Finder grids, which are visible in light microscopy and cryo-ET imaging modalities were used for guidance. Low-resolution images of the grid squares (pixel size 14-28 nm) containing the identified cells or lamellae enabled alignment of light microscopy and cryo-tomography images of the same cell (*37*). Following correlative identification of regions of interest, dose-symmetric tilt series (± 60°, every 3°) were collected in batch mode under minimal dose conditions of about 90-120 e^-^/Å^2^ with defocus between 8 and 14 μm and a magnification resulting in a calibrated pixel size of 0.48 nm.

### Cryo-ET data processing

Automated real-time reconstruction protocols as implemented in the pyCoAn package, version 0.3 (github.com/pyCoAn/distro), an extended python version of the CoAn package (*38*) were employed for all reconstructions. Briefly, immediately after acquisition, tilt series were automatically aligned (*39*) and then reconstructed using the Simultaneous Iterative Reconstruction technique (*40*). Alignment and reconstruction statistics were used to determine quality scores that are provided in real time during data collection. Some of the lamellae tilt series were also reconstructed using EMAN2 (*41*), version 2.9. A total of 387 cellular tomograms were acquired and analyzed. Pseudoviruses were detectable by their distinct structural signature in 181 tomograms. For the analysis of MLV-based particles, 35 tomograms with pseudovirus only, 83 tomograms with hACE2 nanoparticles only and 71 tomograms with a mixture of both were acquired and analyzed. Segmentation of features of interest was achieved using manual tracing with Amira (*42*), version 2019.4. The reconstructions were denoised either with Topaz (*43*), version 0.2.5-2, or non-local means (*44*) followed by a Wiener-like filter accounting for the contrast transfer function at the respective defocus and an estimate of the spectral signal-to-noise ratio (*45*) as implemented in pyCoAn. Tomogram slices were visualized with IMOD (*46*), version 4.11.1, and segmentation results with Amira.

### Sub-tomogram averaging of hACE2 nanoparticles

Sub-tomogram averaging of hACE2 particles on hACE2 nanoparticles was performed following the EMAN2 sub-tomogram averaging pipeline (*47*). Briefly, tilt series from hACE2 nanoparticle samples with high-quality alignment scores and good visibility of protruding particles were binned by two, imported into EMAN2, reconstructed, and the contrast transfer function parameters were estimated for each reconstruction. 311 particles protruding from the membrane and showing good separation from their neighbors were manually selected. After initial alignment and sub-tomogram averaging, the particles were manually curated according to the quality of their fit, leaving 207 particles for the subsequent analysis. These particles were subjected to the sub-tomogram refinement routines in EMAN2, resulting in a final sub-tomogram average of 2.7-nm resolution. The density showed clear evidence for a two-fold symmetry and was symmetrized accordingly for comparison with the hACE2 dimer structure. The hACE2 dimer structure was extracted from the open conformation of the hACE2-B0AT1 complex (*27*) (PDB: 6MD1) and fitted and compared to the density using Chimera (*48*), version 1.15, which was also used to render the images of the density and the atomic model.

## Acknowledgments

We thank Mark H Ginsberg (UCSD) and Edwin Chapman (UW-Madison) for their valuable comments and suggestions on the manuscript; Jean Dubuisson and Muriel Lavie (Institut Pasteur de Lille) for sharing pCMV-MLV-gag-pol and pTG-Luc plasmids for the MLV pseudotyping system and for their continued support and advice; Ignacio Fernández and Félix Rey (Institut Pasteur) for providing soluble, cytoplasmic hACE2 protein samples; Nicolas Escriou (Institut Pasteur) for providing pCI-SARS2-Spike plasmid; Maxime Chazal for providing access to the microplate reader; Pedro Alzari and the Structural Microbiology Unit (Institut Pasteur) for providing access to rotors and chemicals; and Ko-Yung Sit (University of Hong Kong), Michael KY Hsin, and Timmy WK Au (Department of Surgery, Queen Mary Hospital) for their assistance with the *ex-vivo* assays. We acknowledge access to the Leica plunge freezer at the Ultrastructural Bioimaging facility of Institut Pasteur and the Titan Krios, Glacios, Vitrobot, and Aquilos-2 instruments at the NanoImaging Core of the Institut Pasteur.

## Author contributions

DH and NV conceived, directed, and supervised this project. CS, ML, AB, BRdF set up, applied, and refined the methodology, visualization, and analysis. MC, KH, KN and JN performed the *ex-vivo* assays. All authors reviewed and provided input to the manuscript.

## Data and material availability

The data and material generated during the current study are available from the corresponding author upon reasonable request.

## Supplementary Figures

**Fig. S1.**
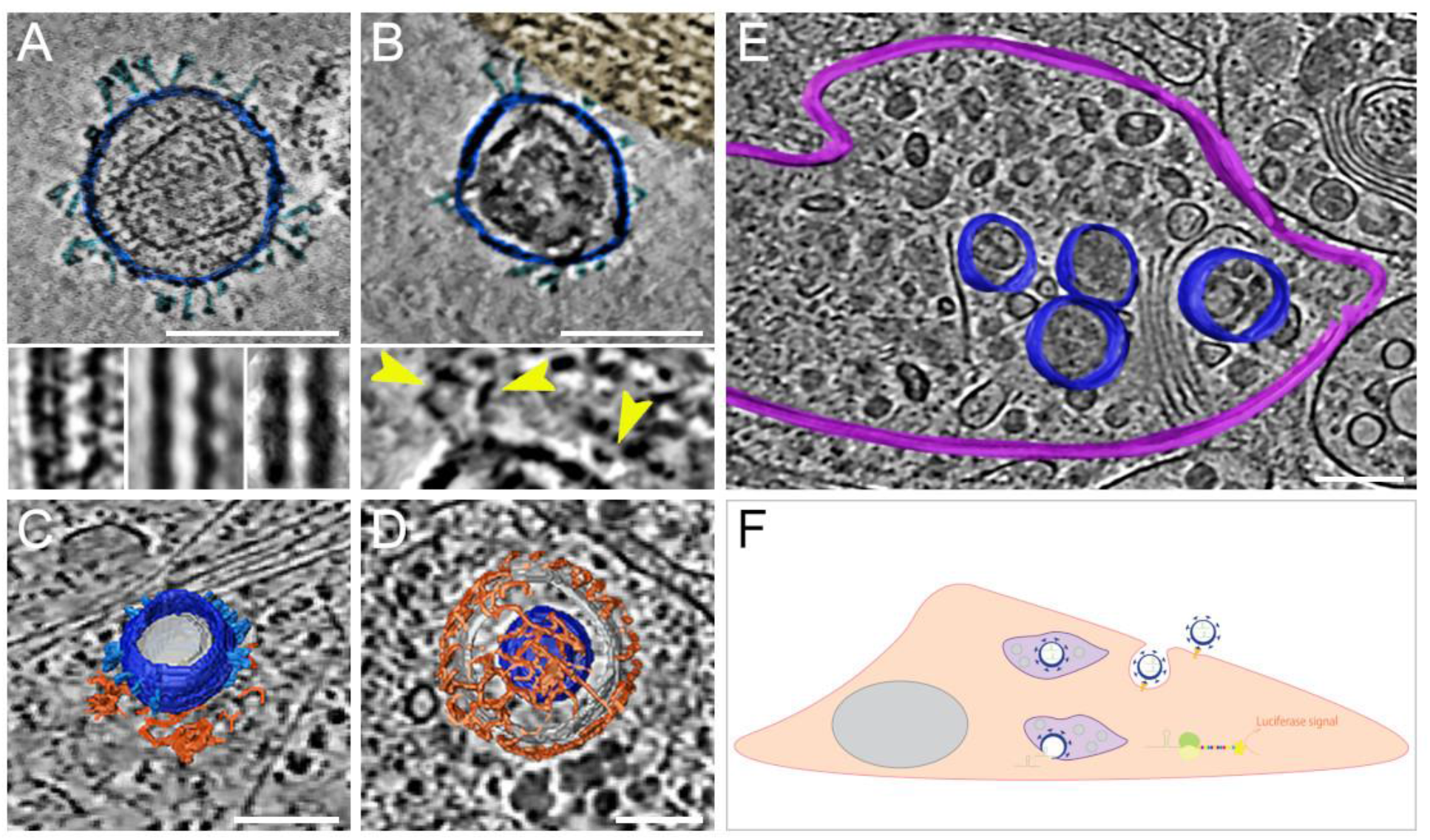
SARS-CoV-2 pseudoviruses enter cells via the clathrin-mediated endocytotic pathway. Cryo-ET reveals pseudovirus morphology and entry mechanism into Vero-E6 cells. Pseudovirus membranes and their segmentation are colored dark blue. Spikes are colored light blue. Clathrin triskeletons are colored in orange, endolysosomal membrane in purple. All virtual slices are 12 nm thick. All scale bars = 100 nm. **(A)** Slice through a tomogram showing a pseudovirus. SARS-CoV-2 spikes are clearly identifiable. The pseudoviruses carry a distinct structural signature imposed by the MLV capsid lattice. The insets show the ridge like structure in a raw slice (left) and after averaging outside (center) and inside (right) the cell. **(B)** Slice through a region of interaction between cell plasma membrane and a pseudovirus. The cell region, tinted gold, shows actin filaments running parallel to the plasma membrane. The inset shows a close-up view with yellow arrowheads pointing at connections between the membrane and the SARS-CoV-2 spikes of the pseudovirus. **(C)** Surface representation of a pseudovirus in the process of entering, attached to a clathrin-coated pit. A slice of the underlying tomogram is superimposed in the background for reference SARS-CoV-2. **(D)** Surface representation of a SARS-CoV-2 pseudovirus inside a clathrin-coated vesicle. A slice of the underlying tomogram is superimposed in the background for reference. **(E)** Surface representation of an endolysosomal compartment containing several pseudoviruses. A slice of the underlying tomogram is shown for reference and represents several loaded vesicles and convoluted membrane structures. **(F)** Schematic representation of SARS-CoV-2-pseudovirus entry and the subsequent steps leading to luciferase signals. Pseudoviruses bind hACE2 receptors via their SARS-CoV-2 spikes on the cell surface (B), which triggers endocytosis (C, D). Once inside, these pseudoviruses are targeted to the endolysosomal system (purple, E) allowing the release of luciferase mRNA. The resulting bioluminescent (yellow star) is proportional to the amount of pseudoviruses that entered the cell.

**Fig. S2.**
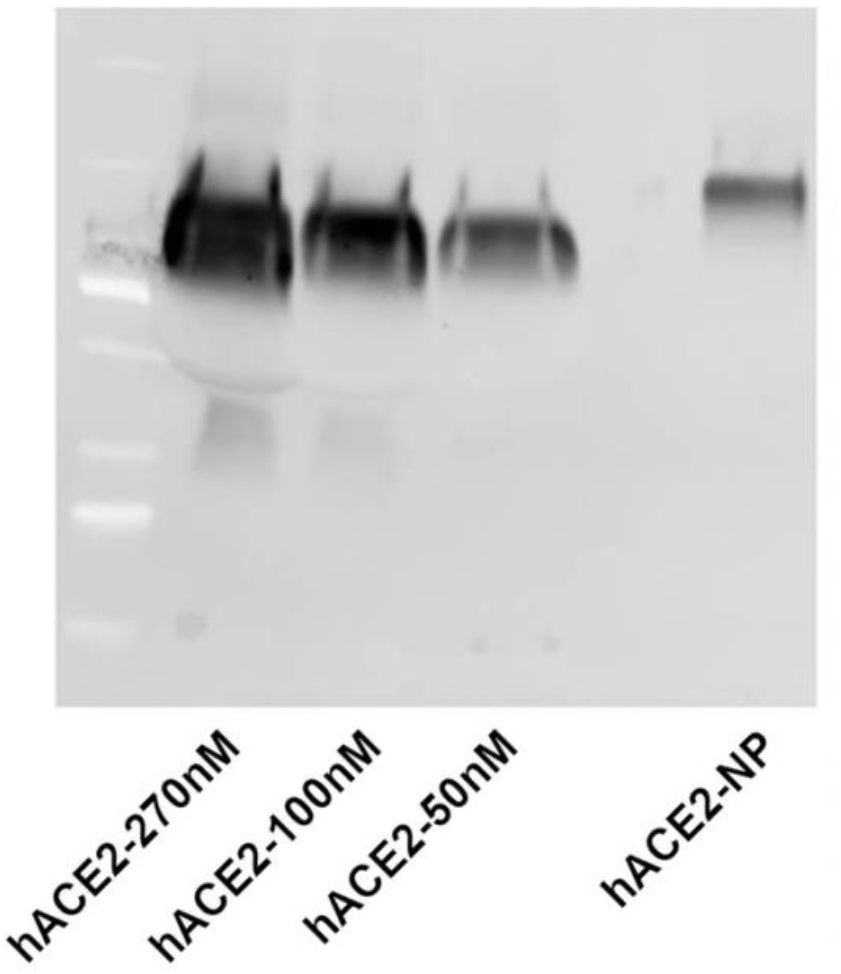
Quantification of hACE2 in nanoparticles. Representative western blot used for quantification of hACE2 levels in hACE2 nanoparticles (hACE2-NP) using soluble hACE2 at different concentrations (hACE2-…nM).

**Fig. S3.**
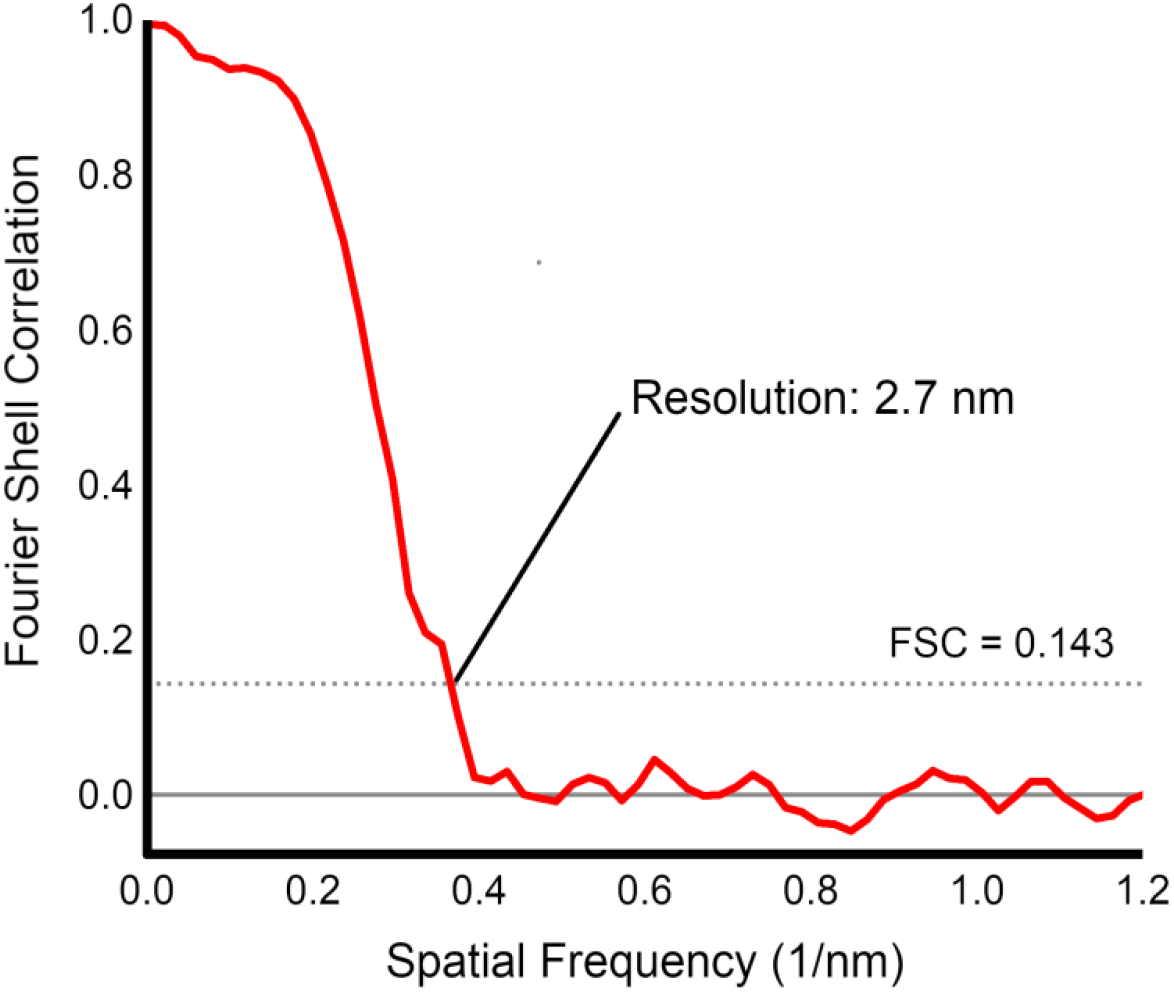
Resolution assessment of nanoparticle-bound hACE2 sub-tomogram average. The data was split in half and refined independently as part of the EMAN2 pipeline. The resolution estimate is then given as the value where the Fourier Shell Correlation (FSC) between the two halves falls below 0.143.

